# Microhabitat use of northern and southern flying squirrels in a recent hybrid zone

**DOI:** 10.1101/2023.05.26.542188

**Authors:** Paul P O’Brien, Jeff Bowman, Sasha Newar, Colin J Garroway

## Abstract

Secondary contact of closely related species may lead to hybridization if reproductive isolation is incomplete. Selection against hybrids may become reinforced if hybrid fitness is low. This can be evident from the divergence of isolating traits. We examined habitat use of northern (*Glaucomys sabrinus* (Shaw, 1801)) and southern (*Glaucomys volans* (Linnaeus, 1758)) flying squirrels in an area of secondary contact in Ontario, Canada. We looked at summer microhabitat use within sites of sympatry and allopatry to test for evidence of reinforcement due to diverging habitat use. We also examined differences in broad-scale habitat features at woodlots to determine predictors of species occurrence across sites. We used 18 years (2002 - 2019) of flying squirrel summer capture data from 6 sites along a north-south transect. Vegetation surveys were conducted at trap locations during summer 2016 to characterize the available microhabitats within sites. Site-level habitat variables were calculated using the collected microhabitat variables. We found microhabitat variables to be weak predictors of trap-level flying squirrel presence and we found no evidence of divergence in microhabitat use over the 18 years. Further, we found latitude, not broad-scale habitat, was the strongest predictor of site-level flying squirrel occurrence. Overall, our findings suggest that microhabitat-based isolation is not being reinforced between flying squirrels; however, hybridization may be limited to areas where climate and habitat is suitable for both species.

## Introduction

Reinforcement is the evolutionary process by which reproductive isolation between diverging species evolves via natural selection in response to maladaptive hybridization (Rundle and Schluter 1998; Noor 1999; Beysard et al. 2015). In areas of secondary contact, interspecific breeding may occur and, if pre-zygotic barriers are not in place, hybrids can be produced. If hybrids are less fit than parental types, then barriers to reproduction, and subsequently assortative mating, are expected to evolve (Servedio and Noor 2003; Matute 2010). Post-mating and post-zygotic barriers on their own still result in wasted reproductive effort (Butlin 1987) thus, pre-mating isolation is required to complete speciation and maximize fitness. The concept of ‘ecological speciation’ is one general model that can explain speciation. It occurs when reproductive isolation evolves as a consequence of divergent selection for traits suited for different environments (Schluter 2001; Rundle and Nosil 2005). When diverging species are allopatric, they cannot produce hybrids and so there can be no direct selection for assortative mating and against interspecies breeding. Thus, areas of secondary contact between recently diverged species are required to study the evolution of reproductive barriers due to reinforcement.

Within the framework of ecological speciation, the most parsimonious explanation for how two coexisting species can arise from a single entity involves an initial phase of geographic isolation and local adaption and a secondary incidence of sympatry where there is reinforcement of reproductive barriers that completes speciation (Grant 2001; Schluter 2001; Rundle and Nosil 2005). Northern (*Glaucomys sabrinus* (Shaw, 1801)) and southern (*Glaucomys volans* (Linnaeus, 1758)) flying squirrels are two species that diverged in allopatry during the early – mid Pleistocene within separate glacial refugia (Arbogast 2007). The contemporary ranges of northern and southern flying squirrels are roughly parapatric, however, due to a more recent (i.e., ∼40 years ago) and rapid range expansion of southern flying squirrels, they have come into secondary contact in Ontario (Bowman et al. 2005). Hybrids have been discovered within a zone of sympatry (Garroway et al. 2010), suggesting that post-zygotic reproductive barriers are incomplete. Secondary contact of flying squirrels within this hybrid zone provides the opportunity to test whether reinforcement will complete the speciation process. Ecological isolation has been shown to play a significant role in maintaining species barriers (Hatfield and Schluter 1999; Wang and Bradburd 2014; Antunes et al. 2021). In such cases, isolation by environment may arise by natural or sexual selection against immigrants, biased dispersal, or low hybrid fitness in parental habitats which reinforces reproductive isolation between parental species (Wang and Bradburd 2014). The distributions of northern and southern flying squirrels are closely associated with boreal and deciduous forests, respectively (Weigl 1968). Therefore, we suggest that reinforcement of ecological isolation between northern and southern flying squirrels is likely to contribute to species barriers in areas of sympatry.

Northern flying squirrels are considered a boreal species with a strong affinity for coniferous and mixed-wood forests. Their strong association with the boreal forest is apparent from a diet largely comprised of lichens and hypogeous fungi (Maser et al. 1985; Li et al. 1986; Trapp et al. 2017). In contrast, the distribution of southern flying squirrels closely follows that of the deciduous, hardwood forests of eastern North America and as such they primarily consume hardwood mast (Sollberger 1940; Muul 1968; Weigl 1978; Helmick et al. 2014). Both species of flying squirrel are considered to be secondary cavity nesters, relying on cavities for feeding sites, safety from predators and weather, and raising young (Weigl 1978; Bendel and Gates 1987). Southern flying squirrels, however, are more reliant on tree cavities, while northern flying squirrels are more flexible in their nest selection, using both external leaf nests and occasionally subterranean nests as well (Trudeau et al. 2011; O’Brien et al. 2021).

The distributions of small mammals are often influenced by associations with macrohabitat features (e.g., food availability, moisture, vegetation type; Bellows et al. 2001; Morris 1979). For northern and southern flying squirrels, divergence in separate glacial refugia has resulted in strong associations with coniferous and deciduous, hardwood forests, respectively. Consequently, their current distributions are strongly tied to these forest types in North America. This is particularly evident at the southern range edge of northern flying squirrels where remnant populations are constrained to high elevations dominated by coniferous species, while southern flying squirrels occupy lower deciduous forests (Weigl 1968; Arbogast 2007). While these sharp vegetational clines can serve to separate closely related or diverging species, macrohabitat associations may be less influential in areas where habitat types (e.g., deciduous and coniferous forests) mix, such as in Ontario, where northern and southern flying squirrels have come into secondary contact and subsequently hybridized (Garroway et al. 2011) as a result of increasingly warmer winters (Bowman et al. 2005). This region is characterized by transitional mixedwood forests (Rowe 1972). Here, more fine-scale selection of habitat or microhabitats within the woodlots used by both species may play a more critical role in separation of the two and subsequent reproductive isolation.

Microhabitats have been defined by Morris (1987) as the environmental variables (i.e., physical and chemical) that influence how an individual allocates its time in space, namely their home range. This can also be classified as fourth-order selection (Johnson 1980). The microhabitat paradigm suggests that sympatry among small mammals is possible through microhabitat partitioning (Jorgensen 2004), a mechanism which has been observed in a number of rodent communities (Morris 1979; Doyle 1987; Magomedov 2021). Within an animal’s home range, not all areas will be used and associations with particular microhabitat features can influence selection of certain areas over others (Karelus et al. 2018). For flying squirrels, like many other animals, habitat features related to daily activities (i.e., foraging and nesting), such as the number of mast or cavity trees, probably make up important microhabitats influencing individual behaviour. Given the known differences in diet and nesting preferences of northern and southern flying squirrels (Dolan and Carter 1977; Wells-Gosling and Heaney 1984), we expect that if reinforcement is acting against hybridization, then microhabitat use related to these differences should play a role.

The objective of this study was to test for evidence of divergence in summer habitat use by northern and southern flying squirrels in an area of close, local sympatry. In Ontario, as part of long-term monitoring of population trends and range boundary dynamics (Bowman *et al*. 2005) we have identified multiple sites where each species occur in allopatry and sympatry. The reinforcement hypothesis predicts that species within an area of sympatry exhibit greater divergence of isolating traits than species in allopatry. Within sites, we hypothesized that if reinforcement is acting against hybridization, then we should see divergence in microhabitat use by northern and southern flying squirrels at sympatric sites through time, thus limiting opportunities for interaction between the two. We predicted that microhabitat variables related to species-specific nest use and diet at sympatric sites would diverge through time, while no divergence would occur at allopatric sites. We also hypothesized that the distribution of flying squirrel species across sites would be predicted by macrohabitat — broad-scale forest characteristics — such that areas of allopatry would reflect habitat preferences of each parent species, while areas of sympatry would be intermediate between the two. Specifically, we predicted that 1) at allopatric northern flying squirrel sites, the occurrence would be positively related to coniferous or coniferous-dominated forests; 2) at allopatric southern flying squirrel sites, occurrence would be positively related to hardwood or hardwood-dominated forests; and 3) sympatry at a site would be promoted by mixed woods.

## Methods

### Study Sites

Study sites were located along a north-south transect from Peterborough, Ontario (44.570°N) to Algonquin Provincial Park (45.584°N; Figure 1). We trapped flying squirrels at 6 sites that represented areas of allopatric northern flying squirrels (*n* = 1), allopatric southern flying squirrels (*n* = 3), and sympatric sites (*n* = 2; Figure 1). All sites used are part of long-term flying squirrel and small mammal research projects (Bowman et al. 2005).

**Figure 1.**
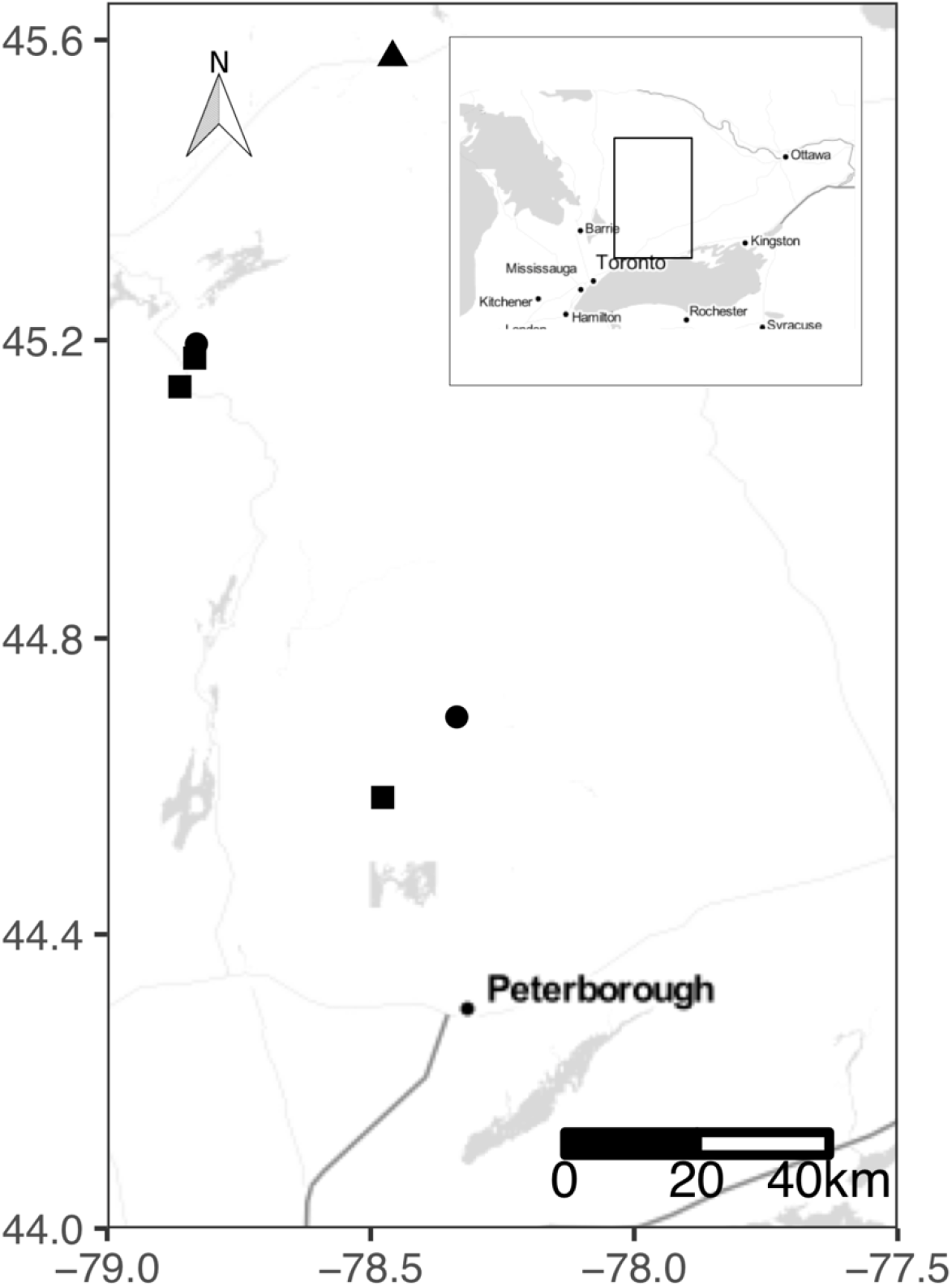
Locations of long-term flying squirrel trapping sites in Ontario, Canada. The 6 sites encompass areas of southern flying squirrel (*Glaucomys volans*) allopatry (square; *n* = 3), northern flying squirrel (*G. sabrinus*) allopatry (triangle; *n* = 1), and sympatry (circle; *n* = 2).

### Species Occurrence

We used 18 years (2002 – 2019) of flying squirrel capture data to estimate species occurrence at trapping sites. Trapping was carried out between June and August of each year for a range of 3 – 7 nights with 20 – 30 traps set per night with about 20 m between traps in either a grid or a transect. The same trap stations were repeatedly sampled throughout the study at each site. We trapped squirrels using Tomahawk model 102 live traps (Tomahawk, WI, USA) baited with sunflower hearts. Traps were mounted on wooden platforms and secured at a height of ∼2m on boles of mature trees and were in place for the duration of the study. If platforms fell down they were remounted and in the rare event of a tree falling over, the platform was replaced on a neighbouring tree. An annual occurrence was recorded for a given trap if a squirrel was ever captured at that trap over the 3-7 night period. All captured individuals were identified to species, sexed, weighed, had age class assigned (adult or juvenile), and were marked with either a 1-g Monel ear tag (National Band and Tag Co., Newport, KY), or a passive integrated transponder (PIT) tags (model TX1411SST, 12.50 mm × 2.07 mm, 134.2 kHz ISO, 0.1020 g; Biomark Inc., Boise, ID, USA) for individual identification, depending on location. Morphological features used to distinguish species included tail and hindfoot length, tail colour, and basal colour of fur on venter (Peterson 1966; Banfield 1987).

### Ethics Approval

All live-trapping and processing methods followed American Society of Mammalogist’s guidelines (Sikes and the Animal Care and Use Committee of the American Society of Mammalogists 2016) and protocols approved by Trent University’s Animal Care Committee (protocol nos. 08034 and 25668).

### Microhabitat Variables

Microhabitat surveys were conducted at each trapping site during the summer of 2016. Within each site, a vegetation survey was conducted within a ∼10m radius of each trap tree as a measure of the available microhabitats at a site. For each survey, species, diameter at breast height (DBH) and decay class (1 = healthy live tree; 9 = decayed stump; (Thomas et al. 1979) of the trap tree were recorded. Researchers followed methodology outlined by Bowman et al. (2001) for vegetation surveys. Within the 10m radius sample plot, all trees ≥8cm DBH were counted, identified to species, and had DBH recorded. Ground cover (i.e., grass, rocks, moss, lichens, leaf, needles, bareground, and shrubs) was measured using a scale of 0 - 5 (0 = 0%, 1 = <1%, 2 = <10%, 3 = 20%, 4 = <50%, 5 = ≥50%) to classify percent coverage. Mast trees, snags, and downed logs within the plot were also counted. The set of microhabitat variables measured is consistent with other studies of flying squirrel microhabitat use (Pyare and Longland 2002; Diggins and Ford 2007; Holloway and Malcolm 2007; Howard et al. 2020). We used the same methods to sample vegetation at trap locations during the summer of 2007. Visual comparison and Pearson correlation tests indicate a strong correlation between the 2007 and 2016 habitat data (Figure S1) and so we chose to use the most recent data from 2016 for the current analysis. In using vegetation surveys from a single year, we assume that temporal variation in microhabitat is consistent across sites. We note that the sites all occurred in unlogged mature forest.

### Site-level Habitat Variables

To estimate site-level habitat variables, we calculated the site-level mean for all continuous microhabitat variables (DBH, canopy coverage, percent lichen coverage, and number of hardwoods, softwoods, mast trees, and snags). For discrete variables (i.e., decay class), we first converted the variable to a binary scale, such that values ≤ 2 were classified as 1 (live) and values > 2 as 0 (dead). We then calculated the proportion of live trees within a site. To further reduce the number of variables, we combined the mean number of hardwoods and softwoods by calculating the proportion of hardwood trees within a site, thus capturing an estimate for both in one variable.

### Mixed Effects Models

To test for evidence of divergence in microhabitat use we ran Bayesian mixed-effects models. For the within-site models, we used either northern or southern flying squirrel occurrence at a trap in a given year as the dependent variable. We selected a subset of microhabitat variables a priori from the total collected that we thought would be most important for flying squirrel habitat use. We chose these variables based on previous research of northern and southern flying squirrels (Bendel and Gates 1987; Pyare and Longland 2002; Diggins and Ford 2007; Holloway and Malcolm 2007; Meyer et al. 2007). To further reduce the number of variables and collinearity, we ran Pearson’s correlation tests. If any two variables had a correlation coefficient ≥ 0.5, we kept the variable we felt was most biologically relevant. The final suite of microhabitat variables we included were: DBH, tree decay, percent canopy, percent coverage of bryophytes, lichens, and logs, and number of softwoods, hardwoods, mast trees, and snags. We also included regional abundance of both species for each year as a covariate to account for variation in occurrence due to natural fluctuations in abundance. Yearly regional abundance was estimated as the total number of each species captured over 100 trap nights. Response variables for within-site models were nested, such that an observation represented the presence of a given species at a trap, within a site for a given year. To capture variation in species occurrence not accounted for by microhabitat or abundance, we included site, year, and trap ID as random intercepts for all models.

For each response (northern or southern flying squirrel occurrence) variable, we fit models for 4 sets of predictor variables including: (1) a null model; (2) variables related to nesting (DBH, decay, canopy, number of hardwoods, number of softwoods, number of snags); (3) variables related to diet (number of logs, bryophyte coverage, lichen coverage, number of mast trees); and (4) a global model, for a total of 8 within-site models. We fit all models in a Bayesian framework with the *brms* package (Bürkner 2017) in R (R Core Team 2020). We ran all models using default, uninformative priors, a warm-up of 5000, and 5000 sampling iterations. For each model we calculated marginal and conditional R^2^ to evaluate and compare model fit using the *performance* package (Lüdecke et al. 2020). Marginal R^2^ describes the variance explained by fixed effects, while conditional R^2^ describes the variance explained by both fixed and random effects.

To test how broad-scale habitat influenced site-level species occurrence, we also ran Bayesian mixed-effects models. For the across-site models, we used the proportion of captures as the dependent variable to capture both species in a single variable. We reduced variables further to include DBH, variance in DBH, percent canopy and lichen coverage, proportion of live trees, proportion hardwoods, and mean number of mast trees and snags. We were interested in testing for divergence in habitat use through time, thus we included year as a random intercept to allow the effect of year to vary between years. We fit two models for the across-site hypotheses: one for habitat variables and the other only including latitude as an independent variable. Models were fit using the *brms* package and R^2^ was calculated with the *performance* package. To help with model convergence, we used weakly informative, flat priors bounded at -10 and 10 to reduce the range of possible parameter estimates explored. As before, we specified 5000 warm-up and sampling iterations.

## Results

### Squirrel Captures

We surveyed 111 trap stations across 6 sites during 2002 to 2019, inclusive. Overall, we had a total of 730 captures over 5511 trap nights. This included 147 captures of 116 northern flying squirrels and 582 captures of 423 southern flying squirrels.

### Within-site Mixed Models

The relationship between most microhabitat variables and squirrel occurrence was not in the predicted direction and 95% credible intervals overlapped 0 (Table 1). For northern flying squirrels, percent lichen coverage and northern flying squirrel abundance were strong predictors of squirrel occurrence for all models (Table 1). According to LOOIC, the diet model was most parsimonious for northern flying squirrels (Table 1). For southern flying squirrels, abundance was the strongest predictor of occurrence for all within-site models, but a weak relationship was found for DBH (Table 2). The nest model was the most parsimonious model for southern flying squirrels according to LOOIC (Table 2). For all within-site models, most variation was captured by the site level random effect (Table 1; Table 2). Site-level effects were in the expected directions for allopatric and sympatric sites (Figure. 2A and B), while random effect of year did not exhibit the expected pattern (Figure. 3A and B).

**Figure 2.**
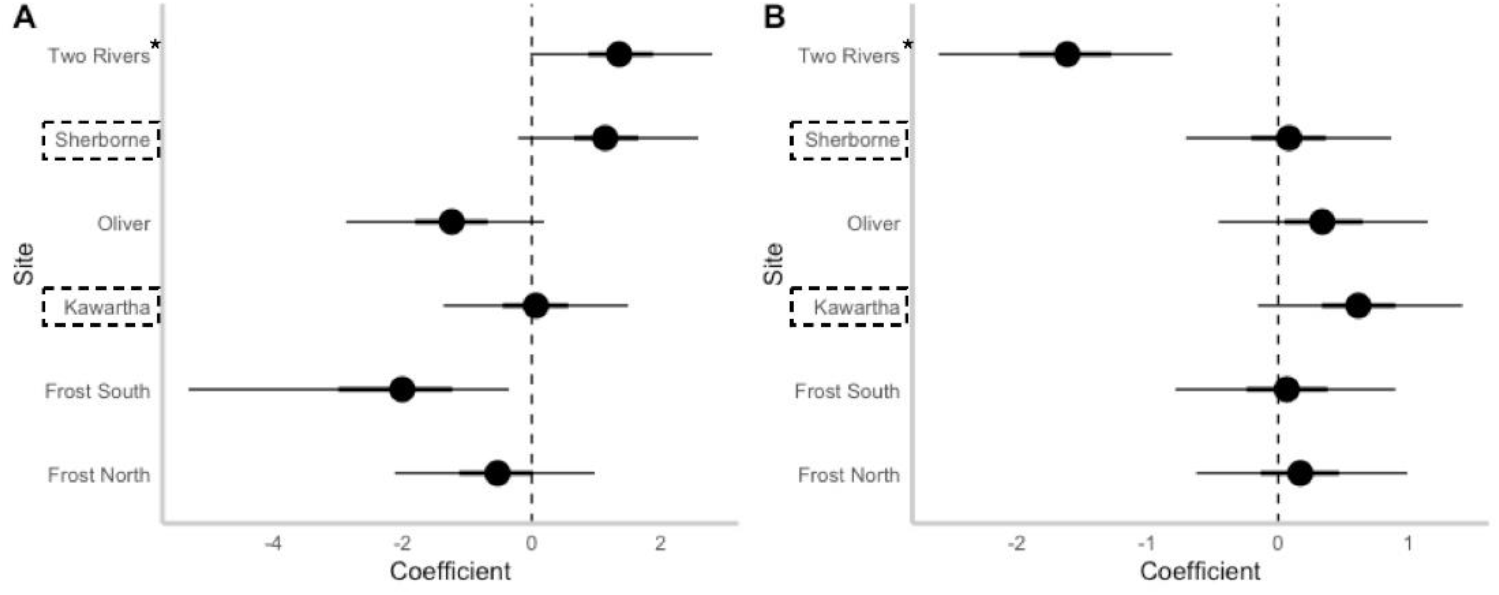
Site-specific parameter estimates for the effect of **A)** diet-related microhabitat variables on within-site northern flying squirrel (*Glaucomys sabrinus*) presence; and **B)** nest-related microhabitat variables on within-site southern flying squirrel (*G. volans*) presence. Ranges are 90% (thick line) and 95% (thin line) credible intervals. Sympatric sites are outlined by dashed boxes, allopatric southern flying squirrel sites are unmarked, and the allopatric northern flying squirrel site is denoted by an asterix.

**Figure 3.**
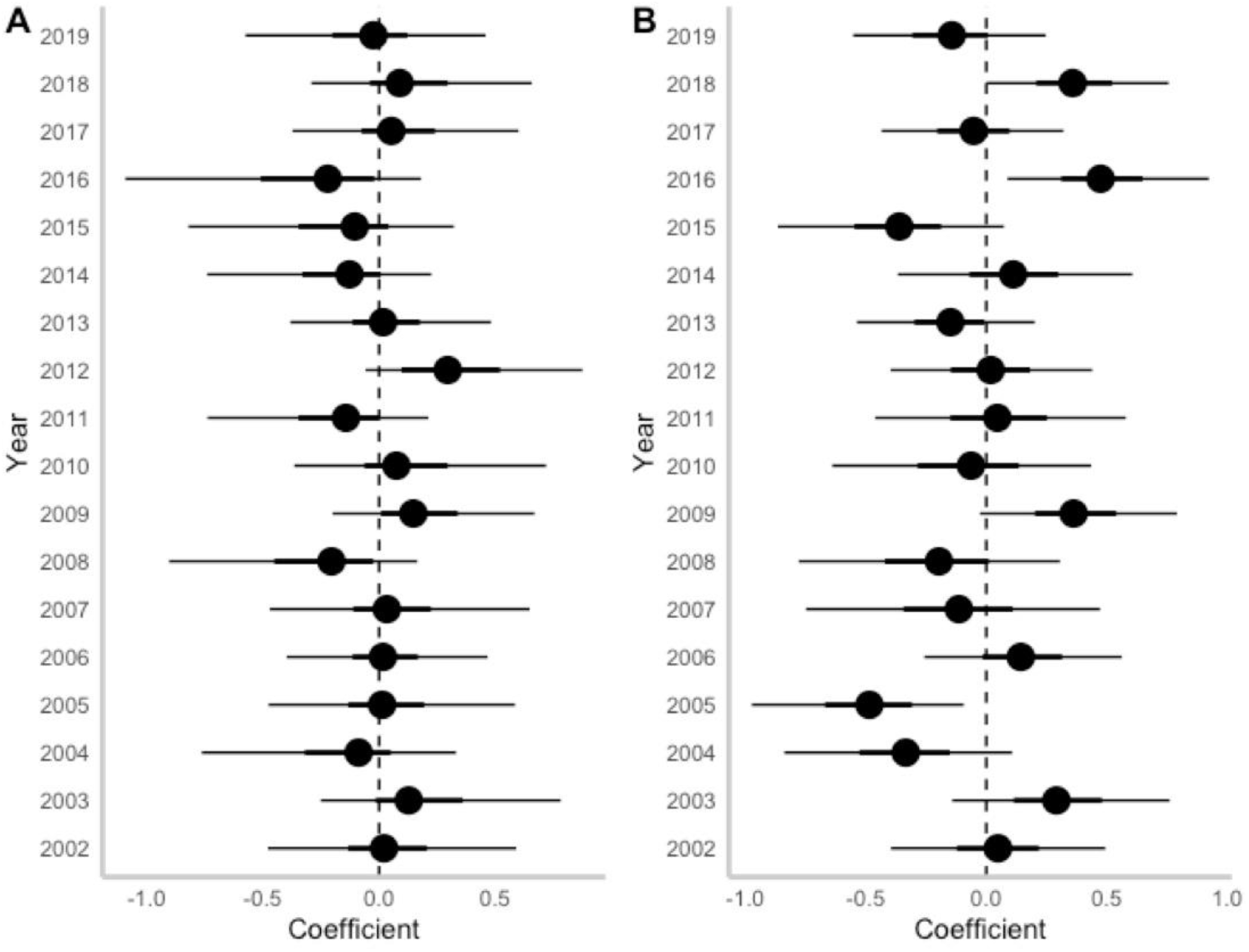
Year-specific parameter estimates for the effect of **A)** diet-related microhabitat variables on within-site northern flying squirrel (*Glaucomys sabrinus*) presence; and **B)** nest-related microhabitat variables on within-site southern flying squirrel (*G. volans*) presence. Ranges are 90% (thick line) and 95% (thin line) credible intervals.

**Table 1.**
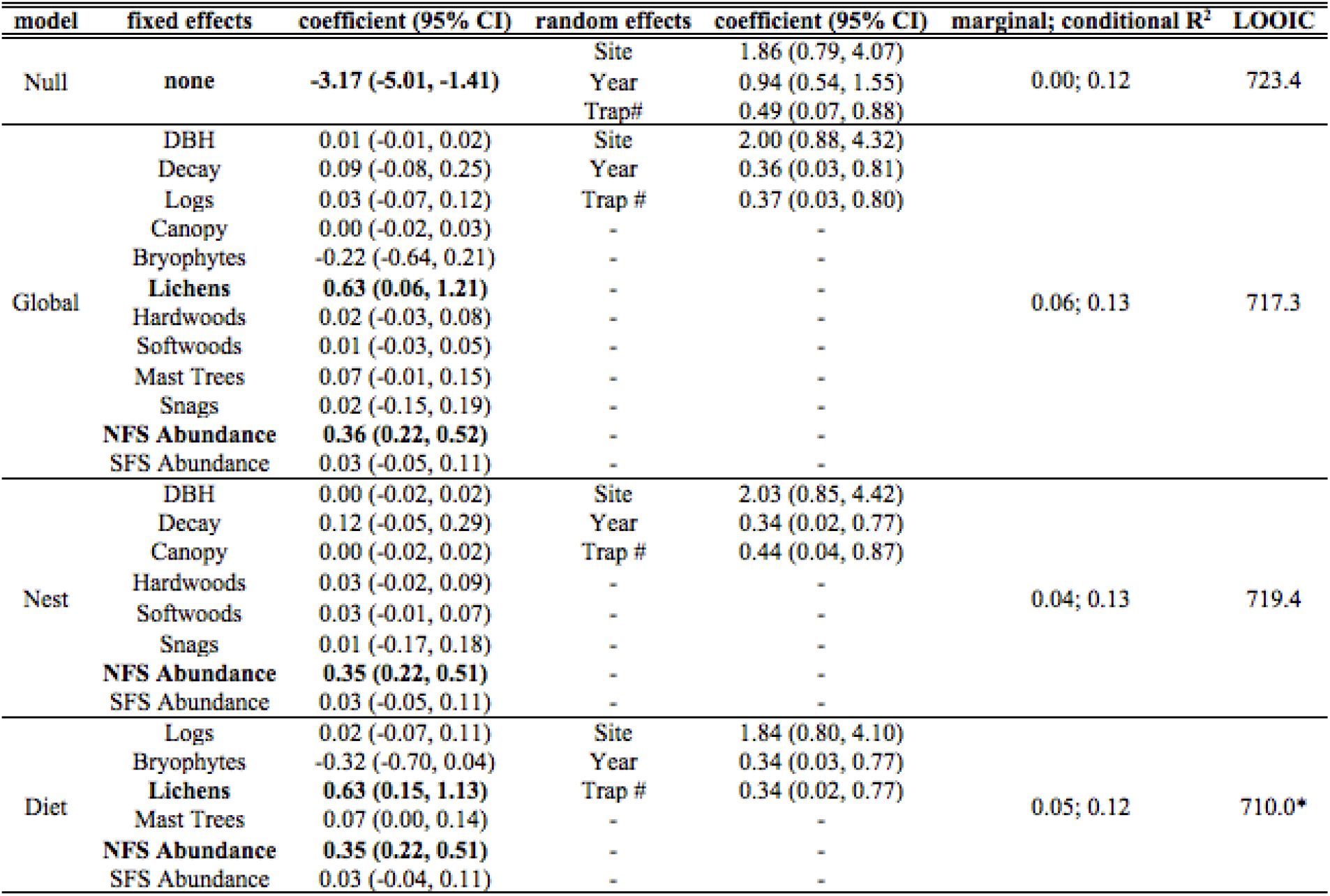
Within-site model summaries for northern flying squirrel (*Glaucomys sabrinus*) trap presence. Four models were fit for the binary response variable each with different sets of predictor variables, including 1) null model, 2) global model, 3) nest model, and 4) diet model. Significant variables are emboldened. Leave-one-out information criterion (LOOIC) indicates model parsimony, with the most parsimonious model indicated by an asterisk.

**Table 2.**
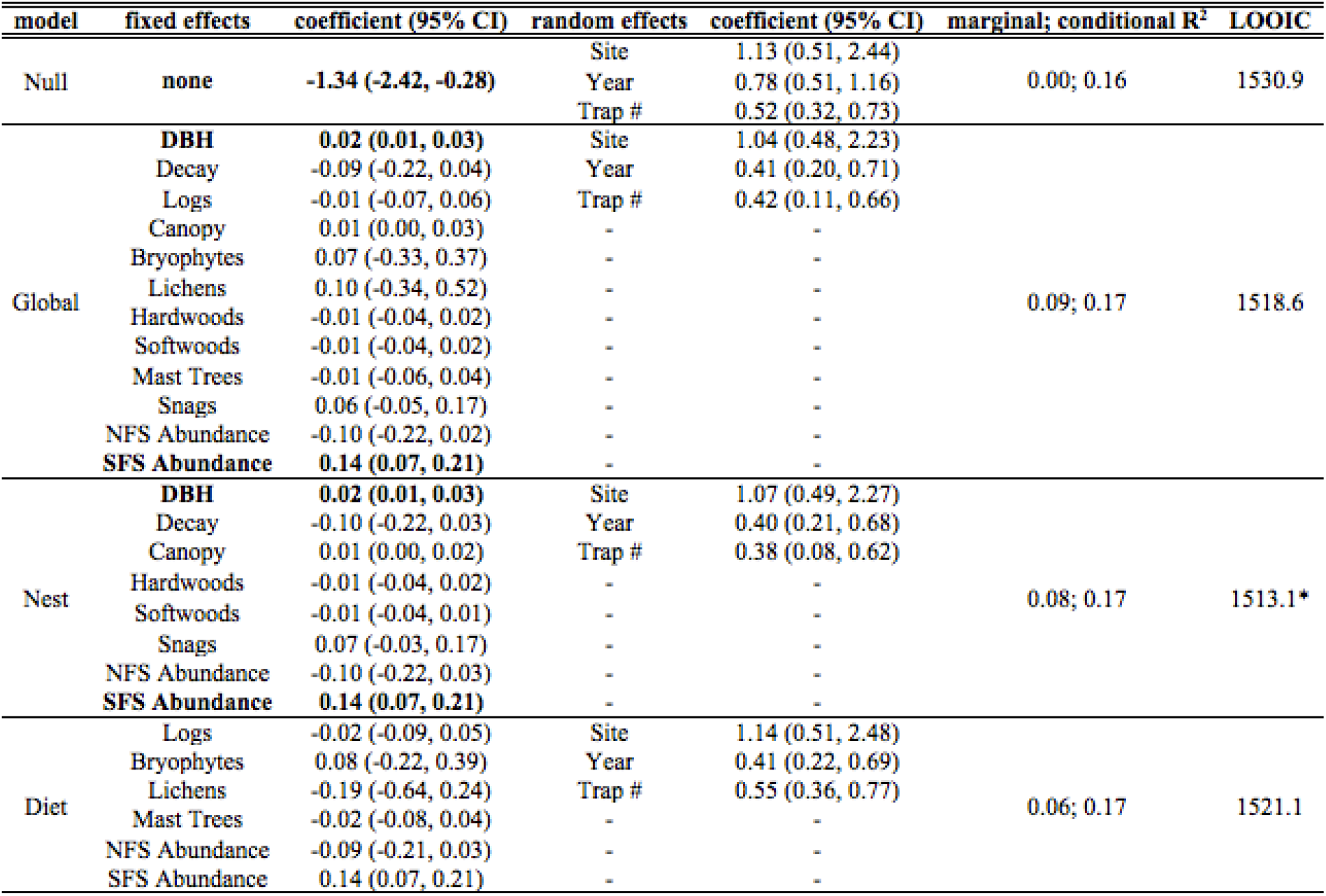
Within-site model summaries for southern flying squirrel (*Glaucomys volans*) trap presence. Four models were fit for the binary response variable each with different sets of predictor variables, including 1) null model, 2) global model, 3) nest model, and 4) diet model. Significant variables are emboldened. Leave-one-out information criterion (LOOIC) indicates model parsimony, with the most parsimonious model indicated by an asterisk.

### Across-site Mixed Models

For site-level models, no relationship was found between any habitat variables and proportion of captures (Table 3). Parameter estimates were not in the predicted direction and 95% credible intervals had wide ranges. The strong positive relationship between proportion of northern flying squirrel captures at a site and latitude was, however, in the predicted direction (Table 3).

**Table 3.**
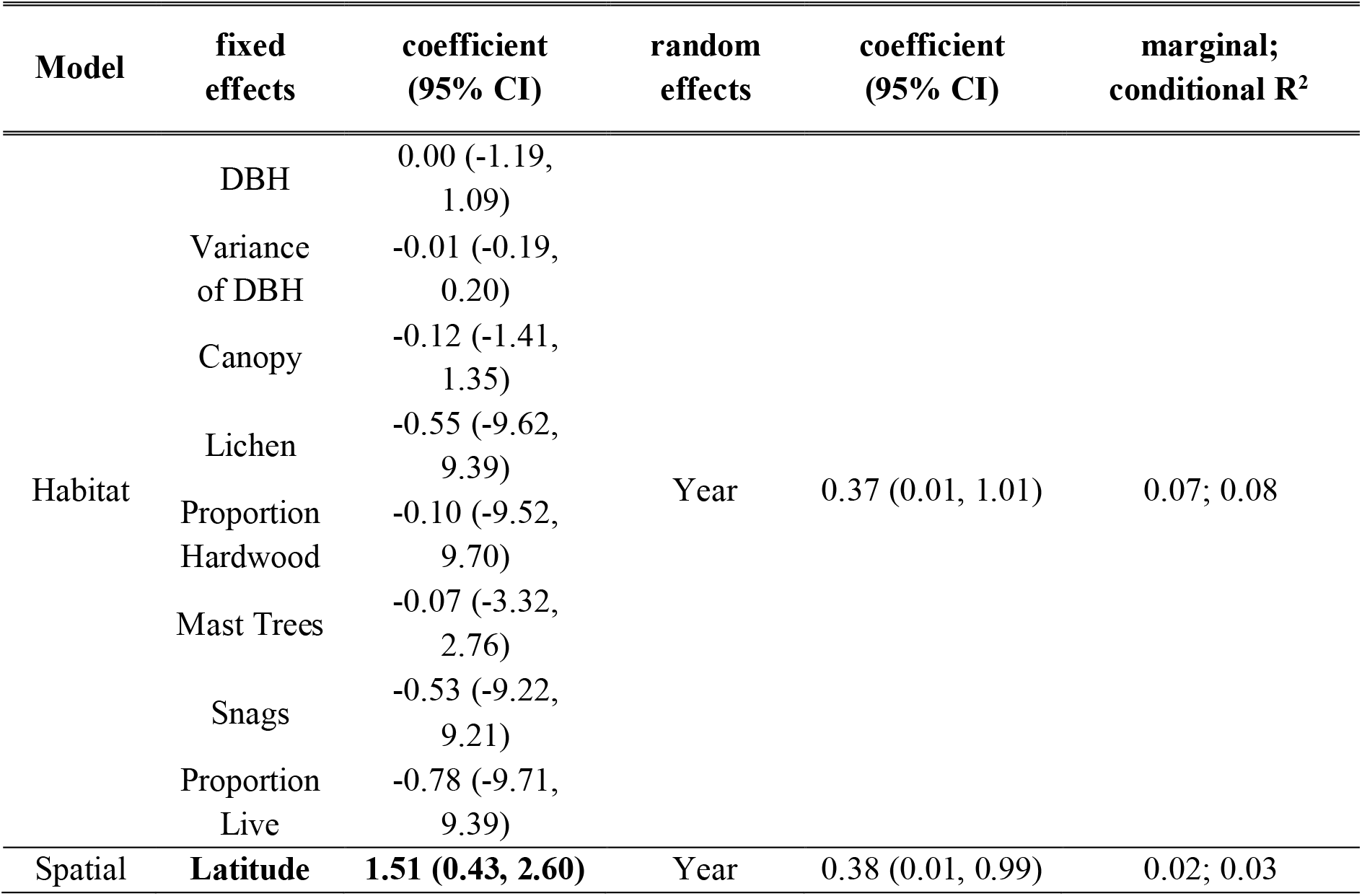
Across-site model summaries for proportion of northern flying squirrels (*Glaucomys sabrinus*) in relation to southern flying squirrels (*G. volans*) captured at a site. Two models were fit for the response variable each with different sets of predictor variables, including 1) a habitat model and 2) a spatial model. Significant variables are emboldened.

## Discussion

In an area of secondary contact between northern and southern flying squirrels and where hybridization has been documented (Garroway et al. 2010), we tested for evidence of divergence in summer microhabitat use. We did not find support for our hypothesis that within sympatric sites the microhabitats used by northern and southern flying squirrels would become more differentiated through time. We also did not find support for our hypothesis that the distribution of flying squirrel species across sites would be predicted by broad-scale forest characteristics, but rather that latitude was the strongest predictor. Overall, our findings do not provide evidence of reinforcement occurring between these two species.

We found the relationship between flying squirrel presence and most microhabitat variables to be weak or not as predicted. Similarly, other studies found northern flying squirrel occurrence to be poorly predicted by many microhabitat variables (Pyare and Longland 2002; Diggins and Ford 2007). We found lichen coverage and regional abundance were the strongest predictors of trap-level occurrence of northern flying squirrels. As would be expected, we found a positive relationship between regional abundance and probability of northern flying squirrel capture. Lichens make up a significant part of the northern flying squirrel’s diet in parts of their distribution (Maser et al. 1985; Trapp et al. 2017), which supports our prediction that microhabitat variables related to diet should influence flying squirrel occurrence. Moreover, it may also be expected that a variable associated with diet would be significant considering the methods used. Live-trapping of flying squirrels occurs at night when squirrels are actively foraging and so where squirrels are captured may be reflective of important foraging areas, thus favouring variables related to diet over those related to nesting. Contrary to this hypothesis, the number of logs surrounding capture sites did not have a strong influence, which is surprising given the association between coarse woody debris and truffles (Amaranthus et al. 1994; Claridge et al. 2000), an important component of northern flying squirrel’s diet (Gomez et al. 2005).

Similarly, the strongest predictor of southern flying squirrel presence was regional abundance. However, in contrast to northern flying squirrels, we found a weak, positive effect of tree diameter (DBH), which is likely explained by nesting preferences of southern flying squirrels as it is an important structural feature in flying squirrel nest selection (Holloway and Malcolm 2007; Zweep et al. 2018; Howard et al. 2020; O’Brien et al. 2021). For both northern and southern flying squirrels, however, the effects of microhabitat variables were weak and most variation in capture probability was explained by random effects, particularly site-level effects.

Despite most variation of within site species presence being explained by site-level random effects, we found no site-level habitat variables to be important predictors of squirrel occurrence at sites. We conclude that this may largely be a result of a low degree of variation in habitat variables among the included sites. Perhaps unsurprisingly, we found latitude to be the strongest predictor of squirrel occurrence among sites. With most of northern and southern flying squirrel’s distributions occurring above and below the 45^th^ parallel, respectively, and our study occurring along a north-south transect (∼ 44.5 – 45.5°N), it makes sense that latitude would explain the greatest amount of variation in occurrence. Historically, the 45^th^ parallel has been considered to be the northern range boundary for southern flying squirrels (Stabb 1988); however, Bowman et al. (2005) found the species range edge to fluctuate in relation to mean seasonal temperatures and mast crop success. That latitude appears to be the strongest predictor of flying squirrel distributions among forest stands would suggest climate is the ultimate driver of site-level occurrence. For the most part, the forest composition of the woodlots used in our study were fairly intermediate between the boreal, coniferous-dominated forests to the north and hardwood forests farther south and so a landscape-level effect of habitat may be more evident at a larger scale where heterogeneity between woodlots would be greater.

We hypothesized that if reinforcement is acting against hybridization, then we should see divergence in microhabitat use by northern and southern flying squirrels at sympatric sites through time. If this was the case, we would predict an increasingly positive year effect, however such a pattern was not observed. A review of small-mammal microhabitat studies by Jorgensen (2004) suggests that most studies fail to detect an effect of microhabitat partitioning among species as a result of inadequate study design. It is possible that we did not include enough microhabitat types in our study, or that trapping effort was low, however, with 6 sites and 18 years of trapping data, we do not think unsubstantial trapping effort is an issue, though we acknowledge our study would have been strengthened with more sampling sites. We note, however, that this long-term study was not designed to test the current question. Moreover, our findings are consistent with numerous other studies using both baited-capture sites and telemetry points that found flying squirrel occurrence (Pyare and Longland 2002; Diggins and Ford 2007) and nest site selection (Holloway and Malcolm 2007) to be poorly predicted by most microhabitat variables. Thus, we suggest microhabitat use may be too fine scale to detect divergence in habitat use within a site for flying squirrels. The scale of effect for such processes may be more evident at a macrohabitat (e.g., home range) level rather than at the individual tree level.

The results of our current study suggest that divergence in microhabitat use between sympatric flying squirrels is not likely occurring; however, differences in nesting ecology and diet may yet play a role in separation of these two species. For example we have observed some weak evidence of divergence in winter nest tree selection (O’Brien et al. 2021). Ecological isolation may be more evident at larger scales, such as at the stand or home range level. During habitat selection experiments, Weigl (1978) found that captive southern flying squirrels chose deciduous habitats markedly when given a choice, while northern flying squirrels were more plastic. However, when the two species were housed together, southern flying squirrels chose deciduous habitat while northern flying squirrels chose coniferous. Weigl (1978) suggested that in areas of sympatry that occur in ecotones between coniferous and deciduous forests, southern flying squirrels may displace northern flying squirrels into coniferous regions. In a system similar to ours, ecologically divergent Douglas (*Tamiasciurus douglasii* (Bachman, 1839)) and red squirrels (*Tamiasciurus hudsonicus* (Erxleben, 1777)) are in secondary contact and hybridize (Fotis et al. 2020). The authors tested for evidence of habitat-based isolation and found that both parental species and hybrids selected similar, but distinct, microhabitats. However, hybrids selected and defended the same preferred macrohabitats as both parental species (Fotis et al. 2020), suggesting that hybrids were not disadvantaged ecologically. In other taxa, macrohabitat partitioning contributes to segregation of closely related species including black-capped (*Poecile atricapillus* (Linnaeus, 1766)) and mountain (*Poecile gambeli* (Ridgway, 1886)) chickadees (Hill and Lein 1988, 1989) and golden-winged (*Vermivora chrysoptera* (Linnaeus, 1766)) and blue-winged (*Vermivora cyanoptera* (Olson & Reveal, 2009)) warblers (King et al. 2015). Therefore, within sites of close local sympatry between northern and southern flying squirrels, such as our study area, divergence in habitat use may be most evident in the habitat types and vegetational characteristics that occur within the home ranges of each species.

In other studies where ecological divergence is clear, selection against hybrids is apparent from reduced fitness of hybrids in parental habitat types (Hatfield and Schluter 1999) or by hybrids only persisting in intermediate habitat zones (Wang et al. 1997). Antunes et al. (2021) found genetic differentiation among subspecies of fire salamanders (*Salamandra salamandra* (Linnaeus, 1758)) to be greatest across sharp transitions of seasonal precipitation and lowest in homogenous regions. We did not examine fitness and putative hybrids were excluded from the current analysis due to low numbers and so we cannot speak to potential fitness reductions of hybrids. However, given that hybrid flying squirrels occur in sympatry with parental species and cases of backcrossing have been documented (Garroway et al. 2010), there appear to be at least some mating opportunities for hybrid squirrels.

From the current findings, it does not appear that northern and southern flying squirrels have diverged in summer microhabitat use through time at our study sites. We suggest that if reinforcement is acting in this area of sympatry, divergence in microhabitat use during the summer is not playing a role in reducing hybridization. Other recent research provides some weak evidence for divergence in winter nest selection (O’Brien et al. 2021), while parasite-mediated competition does not appear to act as a reproductive barrier (O’Brien et al. 2022). Future research should focus more specifically on diet and nesting preferences of the two species and at a spatial scale larger than the microhabitat level. Further, research should also focus on the potential for divergence in winter nesting behaviour since mating occurs in late winter. Species-specific differences in diet and nesting ecology of northern and southern flying squirrels likely contribute to their coexistence in sympatry, however incomplete reproductive barriers still result in occasional cases of non-assortative mating. Further research is required to understand other isolating mechanisms that may be reinforcing species barriers and whether parents and hybrids differ in their survival and fitness. Finally, with a rising awareness of the value of long-term data, systems such as this should continue to be monitored to assess future changes over time.

## Acknowledgements

The authors would like to thank Emily McNaughton, Hannah Hamblin, and Janet Greenhorn for their assistance in the field as well as Mike Brown and Samantha Morin for collection of microhabitat data. The Wildlife Research and Monitoring Section of the Ontario Ministry of Natural Resources and Forestry, the Natural Sciences and Engineering Research Council of Canada, and the University of Manitoba provided financial support. There was no involvement of funding sources in study design, in collection, analysis and interpretation of data, or in the writing of the report.

## Literature Cited

Amaranthus, M., Trappe, J.M., Bednar, L., and Arthur, D. 1994. Hypogeous fungal production in mature Douglas-fir forest fragments and surrounding plantations and its relation to coarse woody debris and animal mycophagy. Can. J. For. Res. 24(11): 2157–2165. doi:10.1139/x94-278.

Antunes, B., Velo-Antón, G., Buckley, D., Pereira, R.J., and Martínez-Solano, I. 2021. Physical and ecological isolation contribute to maintain genetic differentiation between fire salamander subspecies. Heredity 126(5): 776–789. doi:10.1038/s41437-021-00405-0.

Arbogast, B.S. 2007. A brief history of the new world flying squirrels: Phylogeny, biogeography, and conservation genetics. J. Mammal. 88(4): 840–849.

Banfield, A.W.F. 1987. The mammals of Canada. University of Toronto Press, Toronto, ON.

Bellows, A.S., Pagels, J.F., and Mitchell, J.C. 2001. Macrohabitat and Microhabitat Affinities of Small Mammals in a Fragmented Landscape on the Upper Coastal Plain of Virginia. Am. Midl. Nat. 146(2): 345–360. University of Notre Dame.

Bendel, P.R., and Gates, J.E. 1987. Home Range and Microhabitat Partitioning of the Southern Flying Squirrel (Glaucomys volans). J. Mammal. 68(2): 243–255.

Beysard, M., Krebs-Wheaton, R., and Heckel, G. 2015. Tracing reinforcement through asymmetrical partner preference in the European common vole Microtus arvalis. BMC Evol. Biol. 15(1): 1–8. BMC Evolutionary Biology. doi:10.1186/s12862-015-0455-5.

Bowman, J., Forbes, G.J., and Dilworth, T.G. 2001. The spatial component of variation in small-mammal abundance measured at three scales. Can. J. Zool. 79: 137–144.

Bowman, J., Holloway, G.L., Malcolm, J.R., Middel, K.R., and Wilson, P.J. 2005. Northern range boundary dynamics of southern flying squirrels: evidence of an energetic bottleneck. Can. J. Zool. 83(11): 1486–1494. doi:10.1139/z05-144.

Bürkner, P.-C. 2017. brms: A R Package for Bayesian Multilevel Models Using Stan. J. Stat. Softw. 20(1): 1–20. doi:doi:10.18637/jss.v080.i01.

Butlin, R. 1987. Speciation by reinforcement. Trends Ecol. Evol. 2(1): 8–13. doi:10.1016/0169-5347(87)90193-5.

Claridge, A.W., Barry, S.C., Cork, S.J., and Trappe, J.M. 2000. Diversity and habitat relationships of hypogeous fungi. II. Factors influencing the occurrence and number of taxa. Biodivers. Conserv. 9(2): 175–199. doi:10.1023/A:1008962711138.

Diggins, C.A., and Ford, M.W. 2007. Microhabitat Selection of the Virginia Northern Flying Squirrel (Glaucomys sabrinus fuscus Miller) in the Central Appalachians. Northeast. Nat. 24(2): 173–190.

Dolan, P.G., and Carter, D.C. 1977. Glaucomys volans. Mamm. Species (78): 1–6. doi:10.2307/3504026.

Doyle, A.T. 1987. Microhabitat Separation among Sympatric Microtines, Clethrionomys californicus, Microtus oregoni and M. richardsoni. Am. Midl. Nat. 118(2): 258. doi:10.2307/2425783.

Fotis, A.T., Patel, S., and Chavez, A.S. 2020. Habitat-based isolating barriers are not strong in the speciation of ecologically divergent squirrels (*Tamiasciurus douglasii* and *T. hudsonicus*). Behav. Ecol. Sociobiol. 74(3). Behavioral Ecology and Sociobiology. doi:10.1007/s00265-020-2814-5.

Garroway, C.J., Bowman, J., Cascaden, T.J., Holloway, G.L., Mahan, C.G., Malcolm, J.R., Steele, M.A., Turner, G., and Wilson, P.J. 2010. Climate change induced hybridization in flying squirrels. Glob. Change Biol. 16(1): 113–121. doi:10.1111/j.1365-2486.2009.01948.x.

Garroway, C.J., Bowman, J., Holloway, G.L., Malcolm, J.R., and Wilson, P.J. 2011. The genetic signature of rapid range expansion by flying squirrels in response to contemporary climate warming. Glob. Change Biol. 17: 1760–1769. doi:10.1111/j.1365-2486.2010.02384.x.

Gomez, D.M., Anthony, R.G., and Hayes, J.P. 2005. Influence of thinning of douglas-fir forests on population parameters and diet of northern flying squirrels. J. Wildl. Manag. 69(4): 1670–1682. doi:10.2193/0022-541X(2005)69[1670:IOTODF]2.0.CO;2.

Grant, P.R. 2001. Reconstructing the evolution of birds on islands: 100 years of research. Oikos 92(3): 385–403. doi:10.1034/j.1600-0706.2001.920301.x.

Hatfield, T., and Schluter, D. 1999. Ecological speciation in sticklebacks: Environment-dependent hybrid fitness. Evolution 53(3): 866–873.

Helmick, K.R., Barrett, T.L., and Barrett, G.W. 2014. Dietary Resource Preference of the Southern Flying Squirrel (*Glaucomys volans*). Am. Midl. Nat. 171(2): 371–374. doi:10.1674/0003-0031-171.2.371.

Hill, B.G., and Lein, M.R. 1988. Ecological Relations of Sympatric Black-Capped and Mountain Chickadees in Southwestern Alberta. The Condor 90(4): 875–884. doi:10.2307/1368845.

Hill, B.G., and Lein, M.R. 1989. Territory Overlap and Habitat Use of Sympatric Chickadees. The Auk 106(2): 259–268. doi:10.1093/auk/106.2.259.

Holloway, G.L., and Malcolm, J.R. 2007. Nest-tree use by northern and southern flying squirrels in central Ontario. J. Mammal. 88(1): 226–233.

Howard, J.M., Loos, J.E., and Essner, R.L. 2020. Movement and Microhabitat Selection in the Southern Flying Squirrel (Glaucomys volans) in Southwestern Illinois. Northeast. Nat. 27(1): 35. doi:10.1656/045.027.0104.

Johnson, D.H. 1980. The Comparison of Usage and Availability Measurements for Evaluating Resource Preference. Ecology 61(1): 65–71. doi:10.2307/1937156.

Jorgensen, E.E. 2004. Small mammal use of microhabitat reviewd. J. Mammal. 85(3): 531–539.

Karelus, D.L., McCown, J.W., Scheick, B.K., and Oli, M.K. 2018. Microhabitat features influencing habitat use by Florida black bears. Glob. Ecol. Conserv. 13: e00367. doi:10.1016/j.gecco.2017.e00367.

King, R.S., Boysen, J.R., Brenneman, J.M., Cong, R.M., and Hunter, T.S. 2015. Fire created habitat partitioning and isolation between hybridizing warblers. Ecosphere 6(3): art34. doi:10.1890/ES14-00320.1.

Li, C.Y., Maser, C., Maser, Z., and Caldwell, B.A. 1986. Role of three rodents in forest nitrogen fixation in western Oregon: another aspect of mammal-mycorrhizal fungus-tree mutualism. Gt. Basin Nat. 46(3): 411–414.

Lüdecke, D., Makowski, D., Waggoner, P., and Patil, I. 2020. performance: Assessment of regression models performance. CRAN. doi:10.5281/zenodo.3952174.

Magomedov, M.Sh. 2021. Microhabitat partitioning in a rodent community in the arid conditions of the South-western Caspian Lowland. J. Vertebr. Biol. 70(1). doi:10.25225/jvb.20091.

Maser, Z., Maser, C., and Trappe, J.M. 1985. Food habitats of the northern flying squirrel (*Glaucomys sabrinus*) in Oregon. Can. J. Zool. 63: 1084–1088. doi:10.1139/z85-162.

Matute, D.R. 2010. Reinforcement of gametic isolation in drosophila. PLoS Biol. 8(3). doi:10.1371/journal.pbio.1000341.

Meyer, M.D., Kelt, D.A., and North, M.P. 2007. Microhabitat Associations of Northern Flying Squirrels in Burned and Thinned Forest Stands of the Sierra Nevada. Am. Midl. Nat. 157(1): 202–211.

Morris, D.W. 1979. Microhabitat Utilization and Species Distribution of Sympatric Small Mammals in Southwestern Ontario. Am. Midl. Nat. 101(2): 373. doi:10.2307/2424603.

Morris, D.W. 1987. Ecological Scale and Habitat Use. Ecology 68(2): 362–369.

Muul, I. 1968. Behavioral and Physiological Influences on the Distribution of the Flying Squirrel, Glaucomys volans. Univ. Mich. Mus. Zool. Misc. Publ. 134: 1–66.

Noor, M.A.F. 1999. Reinforcement and other consequences of sympatry. Heredity 83: 503–508.

O’Brien, P.P., Bowman, J., Coombs, A., Newar, S., and Garroway, C.J. 2021. Winter nest trees of sympatric northern (*Glaucomys sabrinus*) and southern (*Glaucomys volans*) flying squirrels: a test of reinforcement in a hybrid zone. Can. J. Zool. 99: 859–866. doi:10.1139/cjz-2021-0086.

O’Brien, P.P., Bowman, J., Newar, S.L., and Garroway, C.J. 2022. Testing the parasite-mediated competition hypothesis between sympatric northern and southern flying squirrels. Int. J. Parasitol. Parasites Wildl. 17: 83–90.

Peterson, R.L. 1966. The mammals of eastern Canada. Oxford University Press, Oxford.

Pyare, S., and Longland, W.S. 2002. Interrelationships among northern flying squirrels, truffles, and microhabitat structure in Sierra Nevada old-growth habitat. Can. J. For. Res. 1024: 1016–1024. doi:10.1139/X02-002.

R Core Team. 2020. R: A language and environment for statistical computing. R Foundation for Statistical Computing, Vienna, AustriaR.

Rowe, J.S. 1972. Forest Regions of Canada. Fisheries and Environment Canada, Canadian Forest Service, Ottawa, Ontario.

Rundle, H.D., and Nosil, P. 2005. Ecological speciation. Ecol. Lett. 8(3): 336–352. doi:10.1111/j.1461-0248.2004.00715.x.

Rundle, H.D., and Schluter, D. 1998. Reinforcement of Stickleback Mate Preferences: Sympatry Breeds Contempt. Evolution 52(1): 200–208. doi:10.2307/2410935.

Schluter, D. 2001. Ecology and the origin of species. Trends Ecol. Evol. 16(7): 372–380. doi:10.1016/S0169-5347(01)02198-X.

Servedio, M.R., and Noor, M.A.F. 2003. The Role of Reinforcement in Speciation: Theory and Data. Annu. Rev. Ecol. Evol. Syst. 34(1): 339–364. doi:10.1146/annurev.ecolsys.34.011802.132412.

Sikes, R.S. and the Animal Care and Use Committee of the American Society of Mammalogists. 2016. 2016 Guidelines of the American Society of Mammalogists for the use of wild mammals in research and education. J. Mammal. 97(3): 663–688. doi:10.1093/jmammal/gyw078.

Sollberger, D.E. 1940. Notes on the Life History of the Small Eastern Flying Squirrel. J. Mammal. 21(3): 282–293. doi:10.2307/1374755.

Stabb, M. 1988. COSEWIC status report on the southern flying squirrel *Glaucomys volans* in Canada.

Thomas, J.W., Anderson, R.G., Maser, C., and Bull, E.L. 1979. Snags. In Wildlife Habitats in Managed Forests - the Blue Mountains of Oregon and Washington. Agriculture Handbook No. 553. USDA Forest Service, Washington, D.C. pp. 60–77.

Trapp, S.E., Smith, W.P., and Flaherty, E.A. 2017. Diet and food availability of the Virginia northern flying squirrel (Glaucomys sabrinus fuscus): Implications for dispersal in a fragmented forest. J. Mammal. 98(6): 1688–1696. Allen Press Inc. doi:10.1093/jmammal/gyx115.

Trudeau, C., Imbeau, L., Drapeau, P., and Mazerolle, M.J. 2011. Site Occupancy and Cavity Use by the Northern Flying Squirrel in the Boreal Forest. J. Wildl. Manag. 75(7): 1646– 1656. doi:10.1002/jwmg.224.

Wang, H., Mcarthur, E.D., Sanderson, S.C., Graham, J.H., and Carl, D. 1997. Narrow Hybrid Zone Between Two Subspecies of Big Sagebrush (Artemisia tridentata: Asteraceae). IV. Reciprocal Transplant Experiments Published by: Society for the Study of Evolution Stable URL: http://www.jstor.org/stable/2410963. Evolution 51(1): 95–102.

Wang, I.J., and Bradburd, G.S. 2014. Isolation by environment. Mol. Ecol. 23(23): 5649–5662. doi:10.1111/mec.12938.

Weigl, P.D. 1968. The distribution of the flying squirrels, Glaucomys volans and Glaucomys sabrinus.

Weigl, P.D. 1978. Resource overlap, interspecific interactions and the distribution of the flying squirrels, Glaucomys volans and G. sabrinus. Am. Midl. Nat. 100(1): 83–96.

Wells-Gosling, N., and Heaney, L.R. 1984. Glaucomys sabrinus. Mamm. Species (229): 1–8.

Zweep, J.S., Jacques, C.N., Jenkins, S.E., Klaver, R.W., and Dubay, S.A. 2018. Nest Tree Use by Southern Flying Squirrels in Fragmented Midwestern Landscapes. Wildl. Soc. Bull. 42(3): 430–437. doi:10.1002/wsb.901.

