## Supplemental Table 1 for "Microhabitat use of northern and southern flying squirrels in a recent hybrid zone"

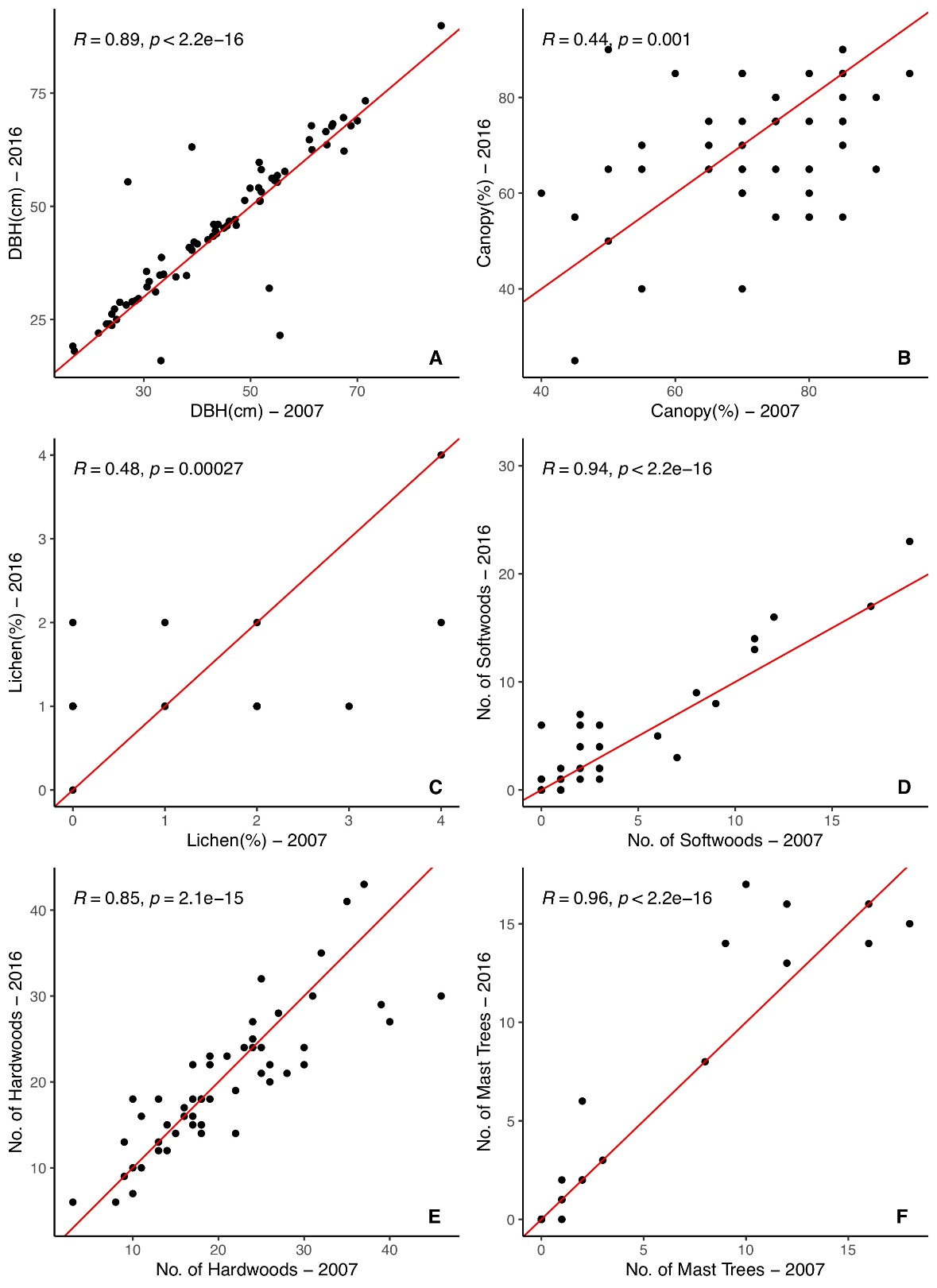

**Figure S1.** Scatterplots and Pearson correlations comparing microhabitat data collected at trap locations from 2016 (data used for current analysis) and 2007 (previously collected data). The red line shows an expected 1-to-1 linear relationship between the recent (2016) and older (2007) habitat data. Data from 2007 was collected following the same methods as described for the 2016 data. The microhabitat variables compared include (A) diameter at breast height (DBH); (B) percent canopy coverage; (C) percent lichen coverage; (D) number of softwoods; (E) number of hardwoods; and (F) number of mast trees.
